# Caloric restriction reprograms adipose tissues in rhesus monkeys

**DOI:** 10.1101/2025.03.03.641286

**Authors:** Josef P Clark, Timothy W Rhoads, Sean J McIlwain, Michael A Polewski, Derek M Pavelec, Ricki J Colman, Rozalyn M Anderson

**Affiliations:** Department of Medicine, University of Wisconsin-Madison, Madison, Wisconsin, United States, 53705; Department of Nutritional Sciences, University of Wisconsin-Madison, United States, 53706; Department of Biostatistics and Medical Informatics, University of Wisconsin-Madison, Madison, Wisconsin, United States, 53792; Carbone Comprehensive Cancer Center, University of Wisconsin-Madison, Madison, Wisconsin, United States, 53792; Department of Animal Sciences, University of Wisconsin-Madison, Madison, WI, USA, 53706; Biotechnology Center, University of Wisconsin-Madison, Madison, Wisconsin, United States, 53706; Wisconsin National Primate Research Center, University of Wisconsin-Madison, Madison, Wisconsin, United States, 53715; Department of Cell and Regenerative Biology, University of Wisconsin-Madison, Madison, Wisconsin, United States, 53705; William S. Middleton Memorial Veterans Hospital, Geriatric Research, Education, and Clinical Center, Madison, Wisconsin, United States, 53705

**Keywords:** adipose, subcutaneous, visceral, caloric restriction, rhesus monkeys

## Abstract

Caloric restriction (CR) is a dietary intervention that delays the onset of age-related diseases and enhances survival in diverse organisms, and although changes in adipose tissues have been implicated in the beneficial effects of CR the molecular details are unknown. Here we show shared and depot-specific adaptations to life-long CR in subcutaneous and visceral adipose depots taken from advanced age male rhesus monkeys. Differential gene expression and pathway analysis identified key differences between the depots in metabolic, immune, and inflammatory pathways. In response to CR, RNA processing and proteostasis-related pathways were enriched in both depots but changes in metabolic, growth, and inflammatory pathways were depot-specific. Commonalities and differences that distinguish adipose depots are shared among monkeys and humans and the response to CR is highly conserved. These data reveal depot-specificity in adipose tissue adaptation that likely reflects differences in function and contribution to age-related disease vulnerability.

**In Brief:** Rhesus monkey adipose tissues are highly responsive to long-term caloric restriction, although the response is depot-specific. Shared features include proteostasis and RNA processing pathways, whereas metabolic pathways, alterations in processes influenced by growth signaling, and changes in exon usage were depot-specific. The impact of CR on subcutaneous adipose tissue is highly conserved between monkeys and humans.

**Highlights:**

- At the transcript level, SAT and VAT taken from the same animals show modest specialization, primarily in metabolic and immune pathways.
- The transcriptome of SAT is more responsive to CR than that of VAT, with depot-specific differences in metabolic adaptation and cell type composition.
- RNA processing pathways are engaged by CR and transcript isoforms are enriched in both depots, but differential exon usage is limited to SAT.
- Adipose reprogramming is linked to CR induced differences in body composition and systemic indices of health.
- The impact of CR on subcutaneous adipose tissue is highly similar between monkeys and humans.

## Introduction

Rhesus monkeys (*Macaca mulatta*) are a highly translational model to study human aging, with genomic, physiological, and behavioral similarities that are shared with humans and a spectrum of age-related diseases and conditions that mirror those prevalent in human aging ^1^. In nonhuman primates (NHP), caloric restriction (CR) without malnutrition improves survival and delays the onset of age-related diseases, disorders, and conditions ^2^. The CR response in NHP involves lower body weight, lower adiposity, lower fasting glucose and insulin, and greater insulin sensitivity ^3^, and is highly similar to that of humans on CR ^4,5^. In mice, CR induces changes in adiposity and in abundance of adipose-derived circulating factors ^6^, pointing to CR-induced differences in adipose tissue mass and adipose tissue function.

Age is associated with changes in body composition where, in general, adiposity increases throughout middle-age, both in humans ^7^ and NHP^8,9^. In general, aging impacts adipose tissue distribution (i.e., where the adipose is in the body) and adipose tissue composition, including changes in adipocyte size, relative proportions of immune and inflammatory cells, vascularization, and fibrosis ^10,11^. Adipose tissue depots are not considered to be equivalent in their contribution to systemic homeostasis and inflammatory tone ^12–14^, with expansion of visceral adipose tissue more often associated with metabolic disease ^15–17^. There is a poorly defined connection between adipose tissue function and whole body inflammatory tone, but aged-related changes in adipose tissue cellular composition and secretory profiles may be a factor in disease vulnerability ^18^.

In terms of total body size and mass, animals on CR are generally smaller and weigh less than Controls with tissue size proportionally reduced; however, the loss is relatively greater for adipose than for other tissues ^19^. There is increasing evidence that adipose tissue from CR animals is also qualitatively different from that of Controls ^20–23^. Adipose tissue function is linked to systemic metabolism, inflammation, and disease ^24,25^. In mice, adipose derived factors responsive to CR such as adipokines ^26^ and extracellular vesicles ^27^ impinge on metabolic and signaling processes in other tissues. In rodents, the reproductive white adipose depot has been better characterized than subcutaneous or visceral depots and as yet the depot-specific impact of CR on adipose remains unclear. Here we describe the transcriptional response of rhesus monkey subcutaneous and visceral adipose depots to CR and document the translatability of these findings by comparison to human data.

## Results

This study uses tissues and data from rhesus monkeys that were part of the Aging and Caloric Restriction study at the Wisconsin National Primate Research Center ^28,29^. A 30% caloric restriction (CR) diet was introduced in adulthood (age 8-14 years) for one half of the monkeys and was maintained throughout the lifespan. As previously reported, CR was associated with improved survival relative to Controls and risk for age-related conditions was significantly reduced ^3^. Samples of subcutaneous adipose tissue (SAT) and visceral adipose tissue (VAT) were collected at necropsy from age-matched (∼25 years of age) adult male rhesus monkeys (n=4 per diet, both depots). For this small cohort, the CR monkeys had significantly lower body weight (∼10kg vs ∼13kg for Controls), fat mass, and lower percentage fat (∼35% vs 18% for Controls), with numerically lower circulating levels of basal insulin and cholesterol, and higher insulin sensitivity (Fig.S1). Glucose levels were not different between diets and were within the healthy range (∼80 mg/dl). Diet did not impact blood cell count (red and white blood cells) and comprehensive blood analyses indicated that there were no differences between Controls and CR for hemoglobin, hematocrit, or for indices of tissue health including kidney (BUN), muscle (creatinine), liver (GTT, ALT, AST) (Fig.S1).

### Transcriptional differences among SAT and VAT are limited to a few genes

The idea that adipose depots show significant differences in function is widely accepted ^14,30^; however, the molecular details are poorly understood. Previous studies in mice did not detect overt baseline transcriptional differences between depots (epididymal v inguinal; ∼3%) ^31^, and similar modest differences have been reported for obese men (visceral v subcutaneous; ∼7%) ^32^. Here we investigated the differences in the transcriptional profiles of SAT and VAT and included the response to a dietary challenge as a means to illuminate differences in functional disposition. Total RNA was extracted from necropsy samples of both adipose depots and libraries were prepared and sequenced yielding an average of 75 million reads with transcripts from 14,749 genes identified across all specimens. Focusing only on Control monkeys, i.e. in the absence of dietary manipulation, the majority (99%) of SAT and VAT genes detected were not differentially expressed (DE) between the two depots (Fig. 1A). DE analysis detected 30 genes with adjusted p<0.05 and 482 genes with unadjusted p<0.05. Among those reaching significance there were several evolutionarily conserved (mice and humans) “depot defining” genes that have established roles in development, including those within the homeobox (HOX) gene network that have been implicated in pattern specification ^31–37^. Genes most divergent in expression between depots include WNT inhibitory factor 1 (WIF1), BMP antagonist gremlin 1 (GREM1), phosphatidyl inositol phosphate kinase (PIK2C2G), and homeobox factors HOXC10, IRX1, IRX6, and NKX3-2 (Fig.1A insets). Adipokine expression ^17,38^ overall was not different between SAT and VAT at the transcriptional level, with only MMP11 reaching significance (adjusted p <0.05), while angiogenesis regulators ANGPT1 and MMP15 were numerically distinct but not significant (Fig.S2). Nor were there overt differences between depots in senescence associated genes documented as part of the SenNet program ^39^ with matrix metalloproteinase MMP3 and chemokine ligands CXCL2 and CCL2 were numerically distinct but not significantly different between the depots (Fig.S2).

**Figure 1:**
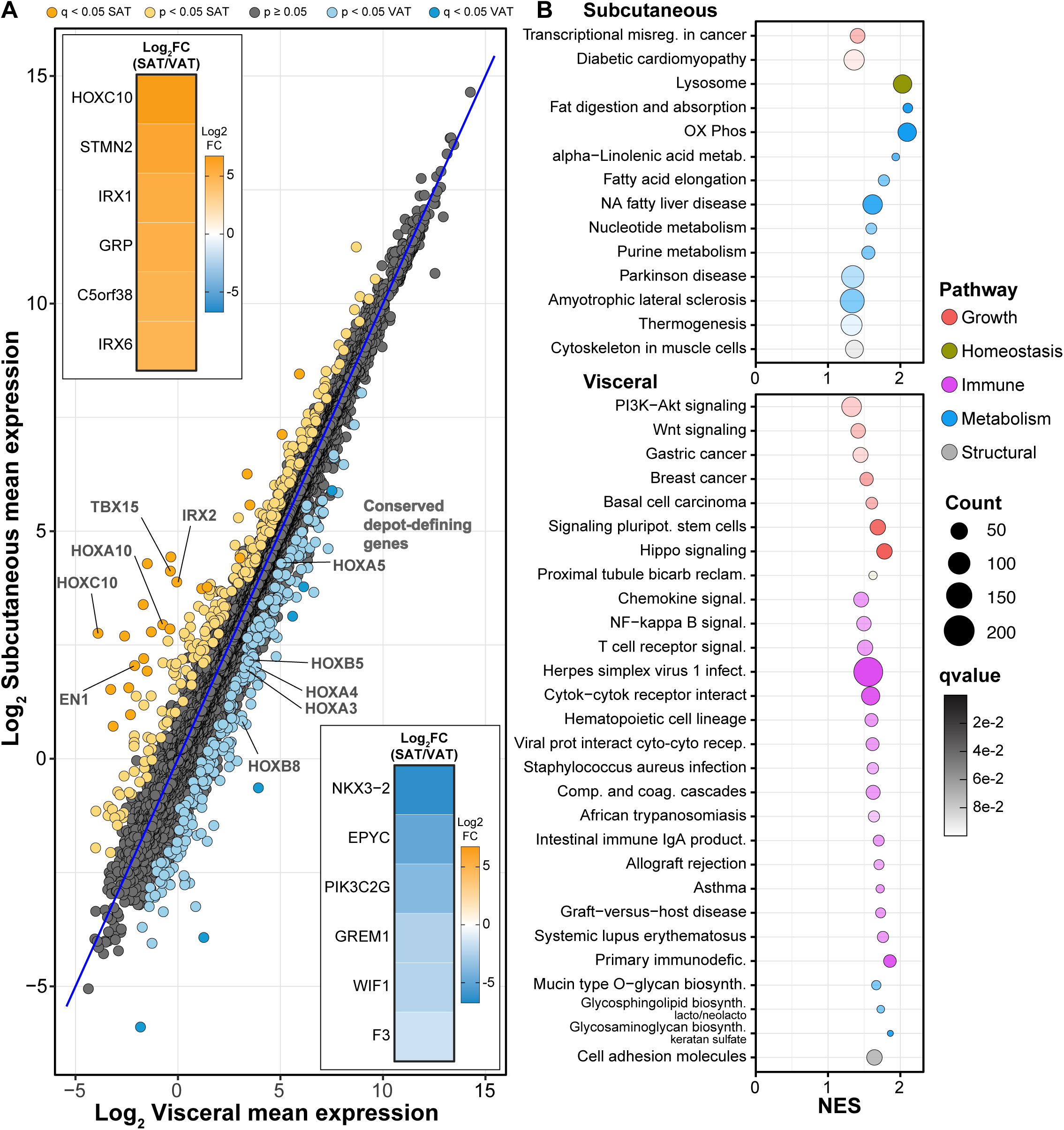
Transcriptional signatures of individual-matched SAT and VAT in rhesus monkeys. (**A**) MA-like plot depicting Log2 mean expression of genes between the two depots in the Control individuals. Genes differentially expressed (adjusted p<0.05) are highlighted for SAT (yellow) and VAT (blue). Inset heat maps depict the top and bottom 20% of DE genes (Log2FC SAT/VAT) for both depots. (**B**) Dot plot depicting KEGG pathway enrichment via GSEA (qvalue<0.1) for SAT (top) and VAT (bottom).

Taking the complete list of all genes represented in either depot, gene set enrichment analysis (GSEA)^40^ with Kyoto encyclopedia of genes and genomes (KEGG) mapping revealed pathways that were enriched primarily in one or the other depot (Fig.1B). The SAT depot-enriched genes were associated with pathways in metabolism and homeostatic processes. The list included the oxidative phosphorylation pathway and pathways associated with lipid and nucleic acid metabolism, indicating that aspects of innate adipose tissue metabolism may be depot-specific. In addition, the lysosome pathway was enriched in SAT relative to VAT. In contrast, VAT-enriched pathways fell largely into categories of growth and immune and inflammatory pathways. Growth signaling pathways including PI3K-AKT, WNT, and HIPPO were VAT enriched. Signatures associated with “inflammaging” ^41^ were prominent in VAT, including cytokine/receptor, innate, adaptive, and autoimmune pathways. Importantly, SAT and VAT depots were taken from the same animals at the same time eliminating the possibility that variability due to differences in systemic factors influenced the adipose tissue transcriptome.

### Transcriptional responses to CR include adipose tissue depot-specificity

In rhesus monkeys, CR is associated with lower adiposity, an outcome that is thought to be linked to improved glucoregulatory function ^1^. In mice, CR reduces the total mass of both SAT and VAT ^42,43^ and the lower adiposity is associated with smaller adipocyte size ^44,45^. CR also impacts factors secreted from adipose tissues ^6,46^, and induces changes in the reproductive adipose tissue transcriptome ^21,47,48^, suggesting that white adipose tissue from CR animals may be qualitatively different from that of Controls. Much of what is known about the impact of CR on white adipose tissue does not compare among depots.

The monkeys in this study were genetically heterogeneous and were at advanced age (∼25 years of age, ∼75 years human equivalent) so that variance among individuals at the transcriptional level was quite high. Accordingly, the impact of CR on the transcriptome of rhesus monkey adipose tissue was modest in terms of statistically significant differences at the individual gene level. The response to CR independent of depot (i.e. depots combined) involved 118 (<1%) DE genes (adjusted p<0.05) and 1322 (9.2%) using unadjusted p values (p<0.05) (Fig.2A). Limiting analysis to SAT only identified 100 significantly DE genes and 834 unadjusted (Fig.2B). For analysis in VAT only, three genes were identified as DE with 637 unadjusted (Fig.2C). The directionality of response was similar between depots for the top DE genes identified in the depot-combined analysis (Fig.2D). CR responsive genes include solute carriers (ion channels: SCN2B, CLCN1, SLC24A2, SCL4A1; metabolite channels: SCL26A7, SLC38A3), and extracellular matrix (MMP3, ADAMDEC1), cell adhesion (FATS2), and cytoskeleton/scaffolding (SPTA1, HEPACAM, GAS2L2) associated factors, and one metabolism associated gene, stearoyl CoA desaturase (SCD1). SAT specific DE genes did not coalesce around any particular pathways but included several development-associated factors (MESP1, WNT5A, WT1, FRZB, TBX1). Taking all DE genes across all comparisons the CR response was generally congruent, although the magnitude of the response differed between depots (Fig.S3). Principal component analysis (PCA) indicates a rough separation of the individual transcriptomes between depots for the CR animals but not for the Controls, indicating that although the depots are quite similar in terms of the transcriptome, the impact of CR is not equivalent between depots (Fig.S3).

**Figure 2:**
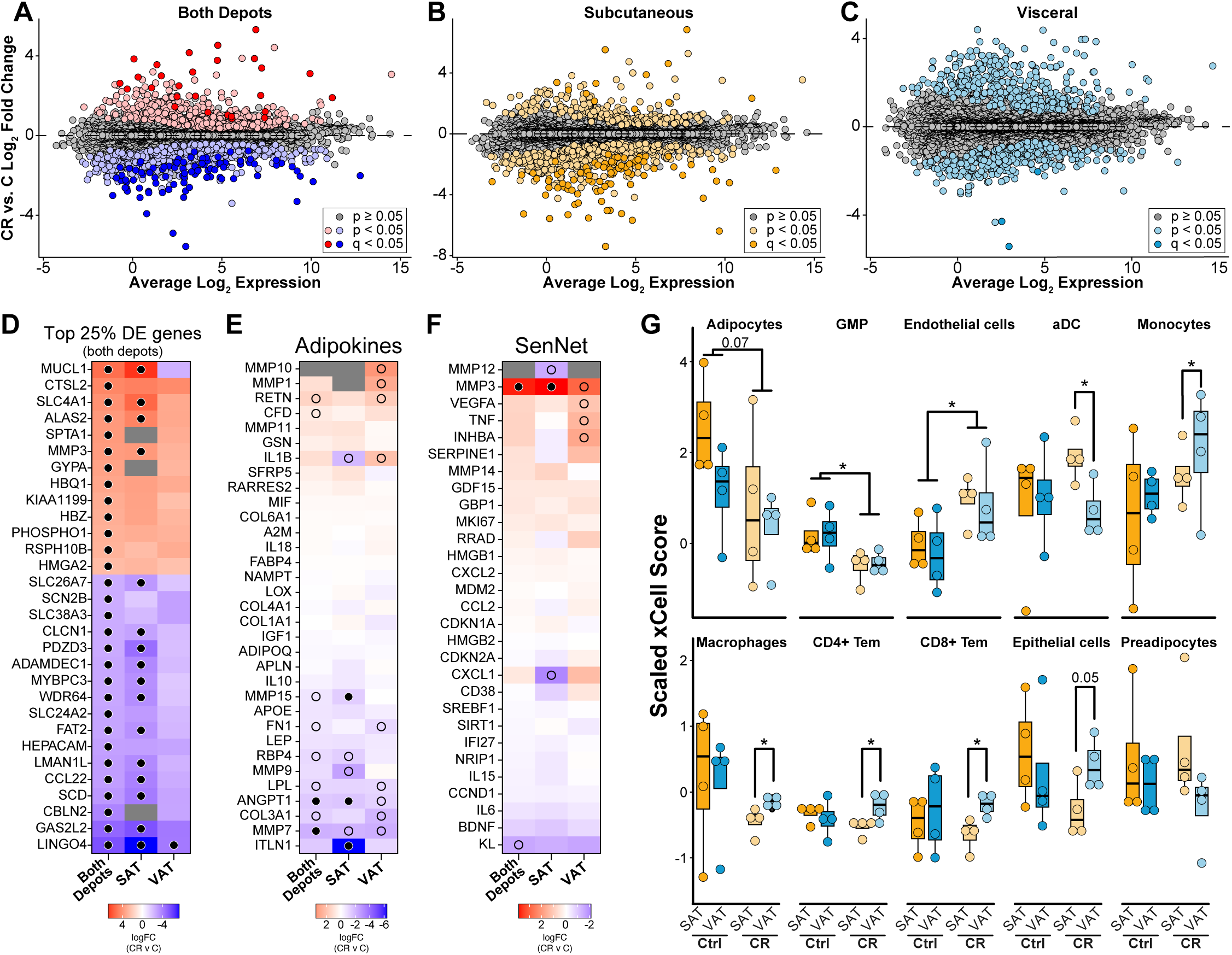
SAT and VAT elicit unique transcriptional responses to CR. MA-plots of genes differentially expressed between CR and Control (C) diets in both depots. Log2 fold change is plotted against the average log2 expression for each gene. Genes differentially expressed are highlighted for each comparison (**A**) both depots (red/blue), (**B**) SAT (yellow), and (**C**) VAT (blue). (**D**) Heatmaps of the CR/C log2FC for the top 25% of most significant DE genes by the absolute value of the log2FC for both depots (shown for each comparison - 30 genes). Heatmaps of the CR/C log2FC for adipokines (**E**) and SenNet factors (**F**). Filled dot denotes passing adjusted p<0.05, empty dot denotes unadjusted p<0.05. (**G**) xCell transcriptome analysis identifying enrichment of individual cell types within each depot on the respective diets (* p<0.05 by Student’s t-test).

### Impact of CR on adipose secreted factors and tissue composition

Adipokine gene expression was somewhat sensitive to CR. In general, CR lowered expression of inflammation associated factors including ECM regulator MMP7 and the secreted glycoprotein ANGPT1 (Fig.2E). Expression of MMP15 and the anti-inflammatory protein omentin (ITLN1) were significantly lower in SAT only (Fig.2F). There were no significant differences in gene expression of the SenNet genes with the exception of MMP3 that was upregulated in SAT only. The relative contribution of different cell types to the captured transcriptome was analyzed via xCell *in-silico* cell-type enrichment ^49^. The cell type distribution was largely similar among depots for Control fed monkeys (Fig.2G), except perhaps adipocytes that had lower representation in VAT than SAT although the difference was not significant. Changes in the representation of adipose immune and inflammatory cells in response to CR have been previously reported in rodents ^50,51^, and in humans on CR ^52^ or during weight loss from obesity ^53^. CR induced significant changes in cell representation where both shared and depot-specific effects were detected. For both depots, CR was associated with relatively lower representation of granulocytes and higher representation of endothelial cells with adipocytes trending lower with CR (Fig.2G). For other cell types the impact of CR was divergent between depots, including the relative representation of dendritic cells, monocytes, macrophages, T cells, and epithelial cells. Together these data show that long-term adaptation to the CR diet in rhesus monkeys involves both shared and depot-specific features, with modest changes in the transcriptome and cell type composition that nonetheless are likely to influence adipose tissue function.

### CR induces common and adipose depot-specific pathways

Two separate approaches were undertaken for pathway analysis: the first combined transcriptomes across depots to establish the adipose tissue “common signature” of CR, and the second took transcriptomes from SAT and VAT independently to establish “depot-specific signatures”. For each approach GSEA via KEGG mapping was used to identify enriched pathways (Fig.3A). Shared pathways identified as significantly enriched with CR in the combined analysis and also significantly enriched in both depots included ribosome and drug metabolism. Other pathways identified as significant in the combined analysis but were significantly enriched in one depot only when the depots were analyzed independently. For example, growth/inflammatory pathways including TNF and JAK/STAT were identified in the combined but were significantly enriched in VAT not SAT. In contrast, spliceosome and proteosome pathways were identified in the combined analysis and were significantly enriched in SAT not VAT. These data suggest that trends in each of these pathways in the “not significantly enriched” depot are sufficient to carry significance when the transcriptomes of both depots are combined. Along these lines, the cell adhesion pathway was identified as negatively enriched with CR in the combined analysis but was not detected as significant in either depot alone. Overall, the SAT CR response was enriched for homeostatic pathways, whereas growth and immune pathways were the principal signatures in the VAT response.

**Figure 3:**
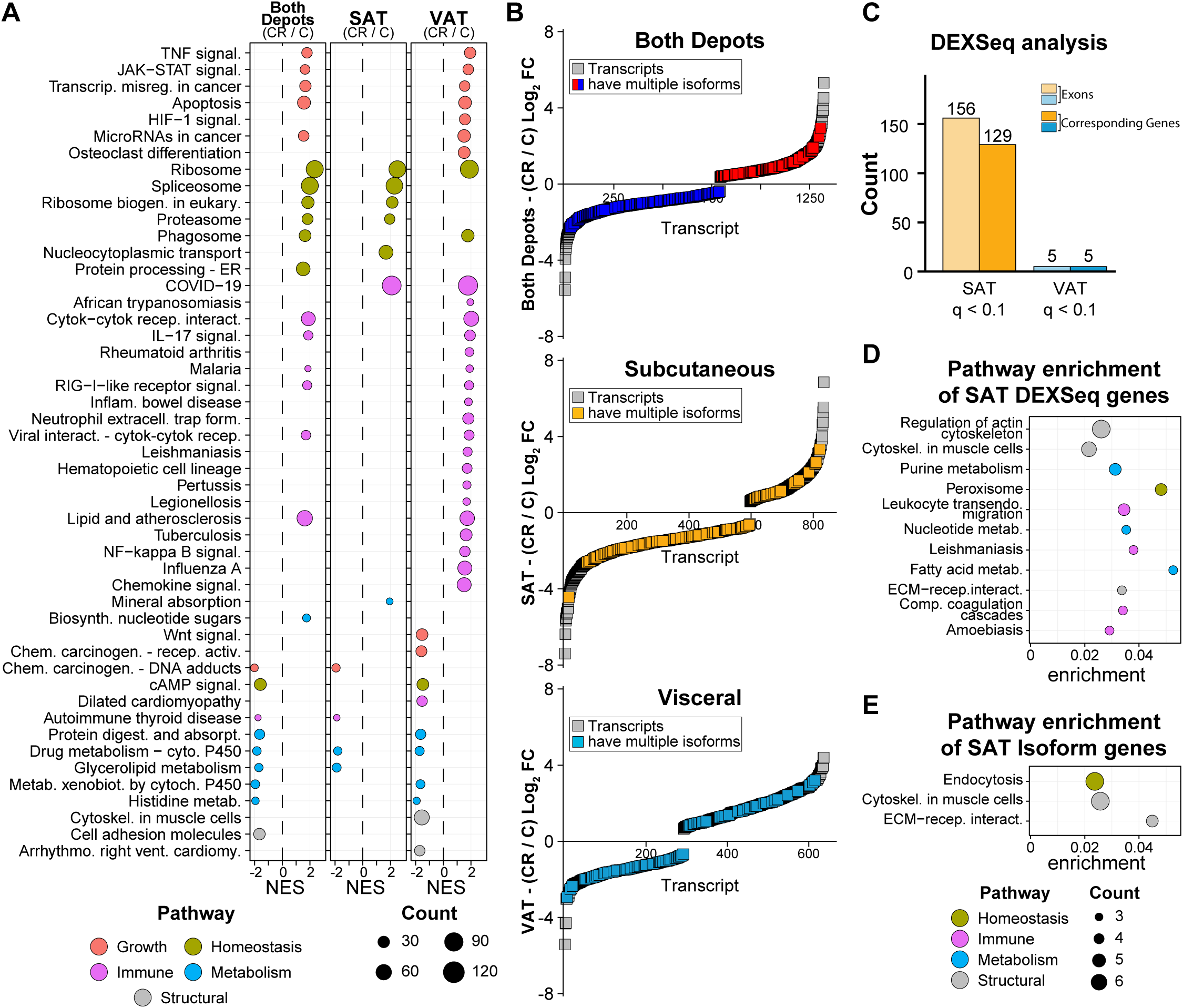
CR induces common and adipose depot-specific pathways. (**A**) Dot plot of KEGG pathways enriched via GSEA (qvalue <0.05) between CR and Control individuals (CR/C) for both depots, SAT, and VAT. (**B**) Rank-order plots of transcript expression by rank against the average log2 fold change between CR and Control diets. Genes where multiple isoforms are detected are denoted for both depots (red/blue), SAT (yellow), and VAT (blue). (**C**) Bar plots showing the number of genes and corresponding exons (passing adjusted p<0.1) that present exon-switching by DEX-seq analysis in response to CR for both SAT and VAT. (**D**) Dot plot of KEGG pathways enriched via ORA (unadjusted p<0.05, gene count > 2) for genes with exon-switching from SAT in (**C**). (**E**) Dot plot of KEGG pathways enriched via ORA (qvalue<0.05) for top-spliced genes (unadjusted p<0.05) in SAT.

### CR engages RNA processing specifically in SAT

Whole transcriptome analysis showed positive enrichment of RNA processing pathways as a common feature of the CR response. Our prior study of the hepatic response to short-term CR in non-human primates identified RNA processing pathways engaged specifically with CR ^54^. Using the “combined” approach with CR responsive transcript isoforms (unadjusted p<0.05), 11% of those genes responding to CR were associated with multiple isoforms (Fig.3B), with roughly the same percentage identified in SAT only or VAT only analysis. To orthogonally measure diet-induced RNA processing, exon usage independent of total transcript levels was quantified using DEXseq with a relaxed significance threshold (adjusted p<0.1). In SAT, 156 exons corresponding to 129 genes that had differential exon usage (Fig.3C). In contrast, in VAT only five exons corresponding to five genes had significantly different exon usage in response to CR. Pathway enrichment using over-representation analysis (ORA) of the genes with differential exon usage in SAT in response to CR identified structural and metabolic processes (Fig.3D). Peroxisome and fatty acid pathways identified here were previously associated with differential exon usage in response to CR, albeit in hepatic tissues rather than adipose tissues ^54^. Similar pathways were identified by analysis of the SAT transcript isoforms associated with the CR response (Fig.3B) using diffSplice, a feature of limma/edgeR that estimates differential exon usage via divergence in sequence read alignments ^55^ (Fig.3E). Cytoskeletal and extracellular structural pathways were the predominant feature identified in SAT from CR monkeys. Based on these data, it seems that the utilization of alternative exons and shifts in splicing might be a conserved mechanism of the NHP response to CR for both SAT and hepatic tissues.

### Integrated adipose tissue response to CR

The monkeys in this study were frequently monitored throughout their lives for biometric, clinical, and disease risk indices (Fig.S1). Multiple factor analysis (MFA)^56^ provides an integrative perspective on health and metabolic indices. Here, sets of variables are structured into groups and their contribution in defining distance among individuals determines their location in the dataspace (Fig.4A), with the strongest features most distant from the origin and related features occupying adjacent space. Fat mass and percent fat were significantly different between Control and CR groups and were clustered together with basal insulin and cholesterol, despite the fact that the latter were not significantly different between groups. The associated principal component analysis separated the individuals based on diet (Fig.4A). To extend and integrate these data, weighted gene co-expression network analysis (WGCNA) ^57^ was used to cluster genes based on expression patterns using transcriptomes from both depots from animals on both diets. WGCNA produced 36 modules (Fig.S4). Module-trait correlations were calculated using the biometric and clinical data (Fig.4B), and the genes within strongly correlated modules were subject to pathway analysis via ORA using the KEGG reference database (Fig.4C). Lean mass and appendicular lean mass were associated with two adipose tissue modules (blue and black) both of which were enriched for metabolic processes. Triglyceride levels were negatively correlated with the adipose tissue module (turquoise) that was enriched for genes involved in homeostatic processes such as proteasome, spliceosome, ribosome, and endoplasmic reticulum associated protein processing. Triglyceride levels were numerically lower in plasma from CR monkeys, although the difference was not significant. Interestingly, triglycerides positively associated with two modules (pale turquoise and blue) that included distinct gene sets but mapped to very similar metabolic pathways.

**Figure 4:**
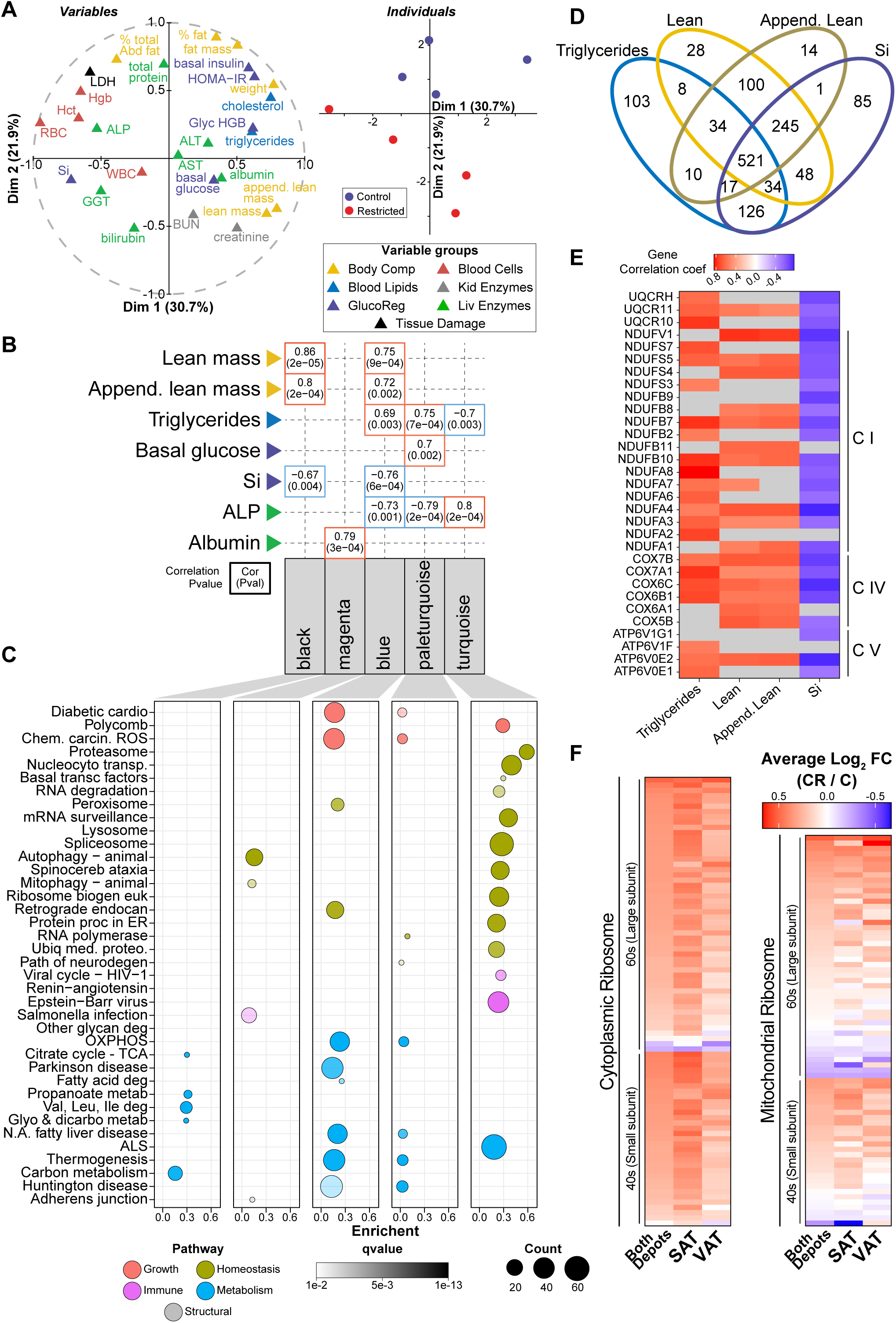
Integrated adipose tissue response to CR. (**A**) Multiple factor analysis of biometric indices: body weight, fat and lean mass, percent fat, appendicular lean mass, percent abdominal fat, HOMA-IR, glycosylated hemoglobin (Glyc HGB), basal glucose, basal insulin, insulin sensitivity (Si), cholesterol, triglycerides, white and red blood cell counts (WBC and RBC), hemoglobin (Hgb), hematocrit (Hct), blood urea nitrogen (BUN), creatinine, lactate dehydrogenase (LDH), gamma-glytamyl transferase (GGT), alanine aminotransferase (ALT), aspartate aminotransferase (AST), alkaline phosphatase (ALP), total protein, albumin, and total bilirubin. Variables correlation plot represents the relationship between each variable and the first two principal components (left), and individuals plot represents the principal component scores for both the Control and CR individuals (right). (**B**) Selected module-trait associations depicting correlations (top number) and pvalue (bottom number). (**C**) Dot plot of KEGG pathways enriched via ORA (qvalue<0.01) for genes contained within each module. (**D**) Venn diagram depicting the overlap of genes significantly associated (p<0.05) with each trait. (**E**) Heatmap of the gene significance (correlation between the gene and the trait) for the KEGG oxidative phosphorylation genes significantly associated (p<0.05) with each trait. (**F**) Heatmap of the Log2FC (CR/C) for the genes contained in the KEGG ribosome pathway for both depots, SAT and VAT.

The association of the blue module with lean mass, appendicular mass, insulin sensitivity, and triglycerides (Fig.S5A) raised questions about the identity of the genes within the module that were driving those associations and whether the same genes were tracking with each of the traits. Within each module, genes highly significantly associated with each trait individually were identified and filtered by significance value (p< 0.05) to determine the extent of overlap among traits (Fig.4D). A group of 521 blue module “core genes” were shared among all four traits and at the pathway level oxidative phosphorylation was common to all four (Fig.S5B). A heatmap of the correlation of individual oxidative phosphorylation gene transcripts in depots combined against trait (Fig.4E) shows some distinction among traits, but in general shows that adipose tissue mitochondrial gene expression is linked to systemic metabolic health (triglycerides and insulin sensitivity) and to animal size (lean mass and appendicular mass).

One of the top pathways responsive to CR for both adipose depots was the Ribosome Pathway. Interestingly, the bulk of cytoplasmic ribosomal genes included in this pathway (69%) were assigned to one WGCNA module (saddlebrown), indicating that the impact of CR on these genes was coordinated. A heatmap of the complete cytoplasmic and mitochondrial Ribosome Pathway gene list reveals strong agreement in the CR response between SAT and VAT (Fig.4F). Data presented here are in agreement with a prior study showing that upregulation of ribosomal genes is part of a conserved transcriptional signature of CR ^58^. Furthermore, these data suggest that ribosomal genes are regulated in concert and that the impact of CR is coordinated among adipose tissue depots.

### NHP adipose tissue profiles and response to CR is conserved in humans

In humans, lower total fat mass is associated with lower comorbidities and a reduced risk of metabolic disease ^59^, but within this association there is a further distinction in that adipose depots are thought to play distinct roles in physiology ^17^. Humans and nonhuman primates are highly similar in adipose distribution and in the impact of age on adiposity ^2,60^. Focusing on the transcriptional level for pathways that are enriched in one or other depot revealed clear parallels between rhesus monkeys and humans (Fig.5A). The publicly available data repository (GEO) was screened to retrieve transcriptional datasets with subcutaneous and visceral depots collected from the same persons, although some studies were conducted in non-healthy individuals. Dual depot adipose tissue data derived from microarray ^61–63^ or from RNA-Seq ^64,65^ were compared against differences detected between depots for the monkeys on this study. In general, the relative enrichment of oxidative phosphorylation, lysosome, and lipid metabolic pathways in SAT compared to VAT was highly similar between monkeys and humans, and the relative enrichment of immune, inflammatory, and cell structural pathways in VAT compared to SAT was also largely conserved.

**Figure 5:**
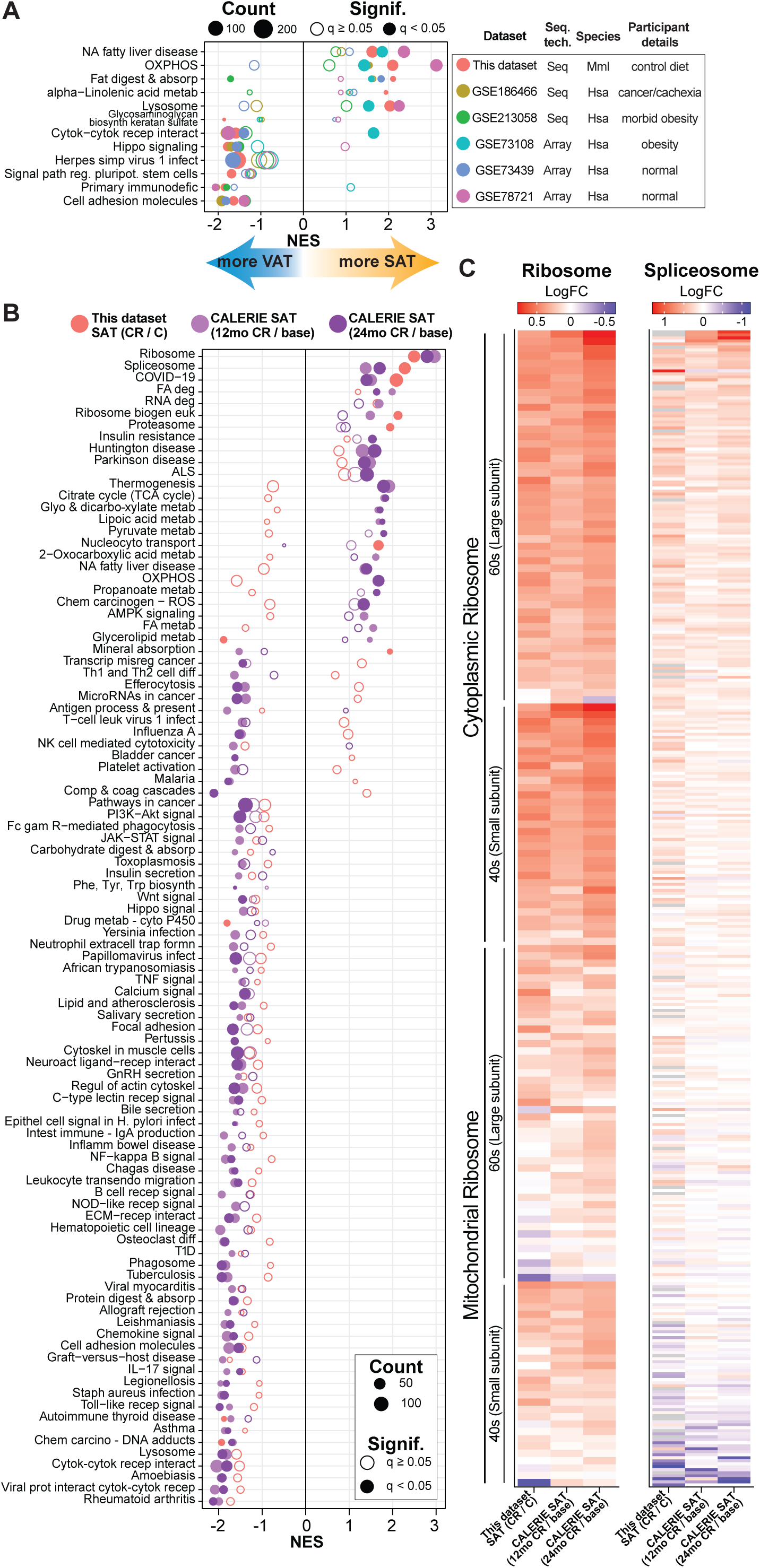
Conservation of adipose tissue profiles and response to CR. (**A**) Dot plot of KEGG pathways enriched via GSEA for NHP (this dataset; SAT/VAT) and human (various GSE datasets; SAT/VAT). The list contains NHP pathways passing qvalue<0.05. (**B**) Dot plot of KEGG pathways enriched via GSEA for NHP SAT (this dataset; CR/C) and human SAT (CALERIE dataset; 12mo/baseline or 24mo/baseline). The list contains at least one comparison with pathway passing qvalue<0.05. (**C**) Heatmap of the Log2FC for the genes contained in the KEGG ribosome and spliceosome pathway for NHP SAT (this dataset; CR/C) and human SAT (CALERIE dataset; 12mo/baseline or 24mo/baseline).

To date, there have been few true caloric restriction studies in humans, with most of the reported studies of lower calorie intake focusing on weight loss in obese individuals. The stand-out study is CALERIE, a three-site two-year intervention study in non-obese adults that showed improvements in multiple risk indices for those on the CR diet ^66–68^. As part of that study SAT samples were taken from a subset of individuals and transcriptional data were generated ^69^. The impact of CR on transcriptional profiles from baseline to 12 months, and 24 months from the human study were compared to the impact of CR on SAT transcriptome in monkeys (Fig.5B). Overall congruence was ∼72% at the pathway level, even though the monkeys were aged and on lifelong CR. Signature positive enrichment of ribosomal, spliceosome, and RNA degradation pathways with CR was aligned between species, as were the negative enrichments in immune, inflammatory, and growth pathways. To explore this further, the impact of CR on individual genes comprising ribosome and spliceosome pathways was quantified for monkeys and for humans at both times points, revealing highly similar patterns of gene expression in the SAT CR response (Fig.5C).

## Discussion

One of the major advantages of the NHP rhesus monkey model is its translatability to human health and aging. Similarities between rhesus monkeys and humans in genetics, physiology, circadian behavior, and age-related disease incidence, are widely accepted ^2,3,70^. Another advantage is the ability to control the environment, diet, and exposures, over the entire lifespan while documenting multiple indices of health longitudinally. The monkeys on this study were part of the WNPRC Aging and Calorie Restriction study that involved longitudinal assessments of health status from adulthood ^28^, so that biometric, blood chemistry, and glucoregulatory status in the months prior death were documented. WGCNA analysis identified adipose transcriptional signatures that correlated with whole body and systemic indices, notably the connection between adipose tissue metabolic pathways and lean mass, insulin sensitivity, and circulating triglyceride burden. These data suggest that adipose tissue is sensitive to differences in body composition and that the metabolic status of adipose tissue is reflected in differences in circulating factors associated with disease risk. The relative contributions of SAT and VAT to systemic outcomes could not be established using this approach because the biometrics and circulating factors are mapped to each individual monkey and the depots are not considered independently.

In this study, the emphasis was on directly comparing the impact of CR across two adipose depots. As previously reported, survival is extended in CR animals; however, the individual monkeys in this study were selected to be age-matched to eliminate age as a confounder and were not different in terms of survival. Of the Control animals two of four died of age-related conditions (cancer, renal and cardiac disease), and all remaining monkeys died of non-age-related conditions (NARDs) ^3^. Although matched for chronological age, the biometric, blood chemistry, and glucoregulatory function data suggest that CR was beneficial in terms of physiology and systemic indices of disease risk. It seems reasonable to conclude that the CR intervention was effective in delaying biological aging of the monkeys in this current study.

The alignment in depot-specific transcriptional signatures of SAT and VAT between monkeys and humans points to innate and conserved differences in adipose tissue function dependent on location in the body. In general, the transcriptome for both depots was highly similar with surprisingly few genes identified as DE. It seems likely that modest differences in expression among other transcripts are the basis for the differences in pathway enrichment between depots. The relative enrichment in metabolic pathways in SAT and immune and inflammatory pathways in VAT reported here aligns with clinical studies that have identified VAT as the “bad fat” ^71,72^. In obese humans, acute low-calorie diet weight loss is associated with a reduction in VAT but not SAT depot size whereas a chronic low-calorie diet seems to impact VAT and SAT ^73,74^. Interestingly, in this long-term CR intervention study a greater number of DE genes were identified in SAT than in VAT, with clear differences in the pathways associated with the CR response for the two depots.

Two major pathways identified in this study as being responsive to CR in adipose tissues include RNA processing and ribosomal pathways. For the former, increasing evidence suggests that CR engages RNA-processing pathways in other tissues including liver ^54^ and muscle ^75^ in NHP, and in other species including humans, mice, and worms ^76–78^. We find that the CR involves alternative isoform transcripts in both depots, but exon utilization is engaged primarily in SAT. Ribosomal genes are notably upregulated in both depots in response to CR. Transcriptome analysis of liver tissue from NHP following a 2-year transition onto a CR diet also show that the ribosome pathway is upregulated ^54^. Similarly, ribosome pathways are upregulated in the liver of mice on long-term CR and in genetic models of longevity ^79^. Here the broad impact of CR across both small and large ribosomal units was conserved between monkeys and humans. The physiological consequence in terms of ribosomal composition, ribosomal function, and translation selectivity is still unclear. A surprising outcome of the study was the lack of transcriptional signatures aligned with adiposity, a phenotype that was significantly different between Control and CR fed animals. Not only were there no modules of genes associated with adiposity by WCGNA, but targeted regression analysis also failed to identify signatures reflecting this key difference in body composition. Perhaps a change in adipose function rather than adipose mass is a driving feature in aging. Function may be altered independent of changes in adiposity or perhaps changes in the adipose transcriptome in response to depot size are distributed across processes. Either way genes correlating with differences in adiposity did not coalesce into distinct pathways.

Limitations of the study: The number of monkeys (n=4 per diet group) is on the lower end; however, the fact that both depots were taken from the same animals for which there are extensive biometrics and clinical measures offsets this concern somewhat. The comparisons made of transcriptional data from this study and that derived from independent human studies adds weight to the findings and indicates that at least certain aspects of adipose tissue depot specialization and the response to CR reported here for NHP is conserved in humans. Nonetheless, the findings from this study should be considered preliminary. A second limitation is that this study was conducted using males only. The monkeys were from the first group enrolled in the larger cohort and were selected to match for age at death. The choice was motivated by the possibility that changes with age are not linear and that differences in life-stage would introduce confounding effects. This strategy prevented any insight into potential sex dimorphism in the response of adipose tissues to CR and limited insights gleaned to the impact of CR in aged male animals. The main cohort includes both males and females with median age at death of 26.4 years (IQR: 7.2 years) that will allow an investigation of the impact of age, sex, and the CR diet.

Overall, this study supports the hypothesis of a conserved transcriptional program in the CR response. Parallels in the impact of CR on RNA processing and ribosomal genes for adipose (this study), liver ^54^, and skeletal muscle ^75^ argue that these are core features in CR’s mechanisms, but the physiological consequence of these common adaptations remains unclear. The congruence between humans and monkeys in response to CR suggests that the health benefits are linked to changes in adipose but how this plays out in terms of survival remains to be seen. The concept of adipose tissue as a driver in setting metabolic status in the context of aging, health, and disease has gained traction ^18,80^. At later ages, survival is linked to the retention of adipose tissues, and severe loss of adipose is associated with increased risk of mortality ^81^. It will be important to establish the extent to which adipose tissue directs the pace of aging and how that influences health status into advanced age.

## Materials and Methods

### Key Resources Table

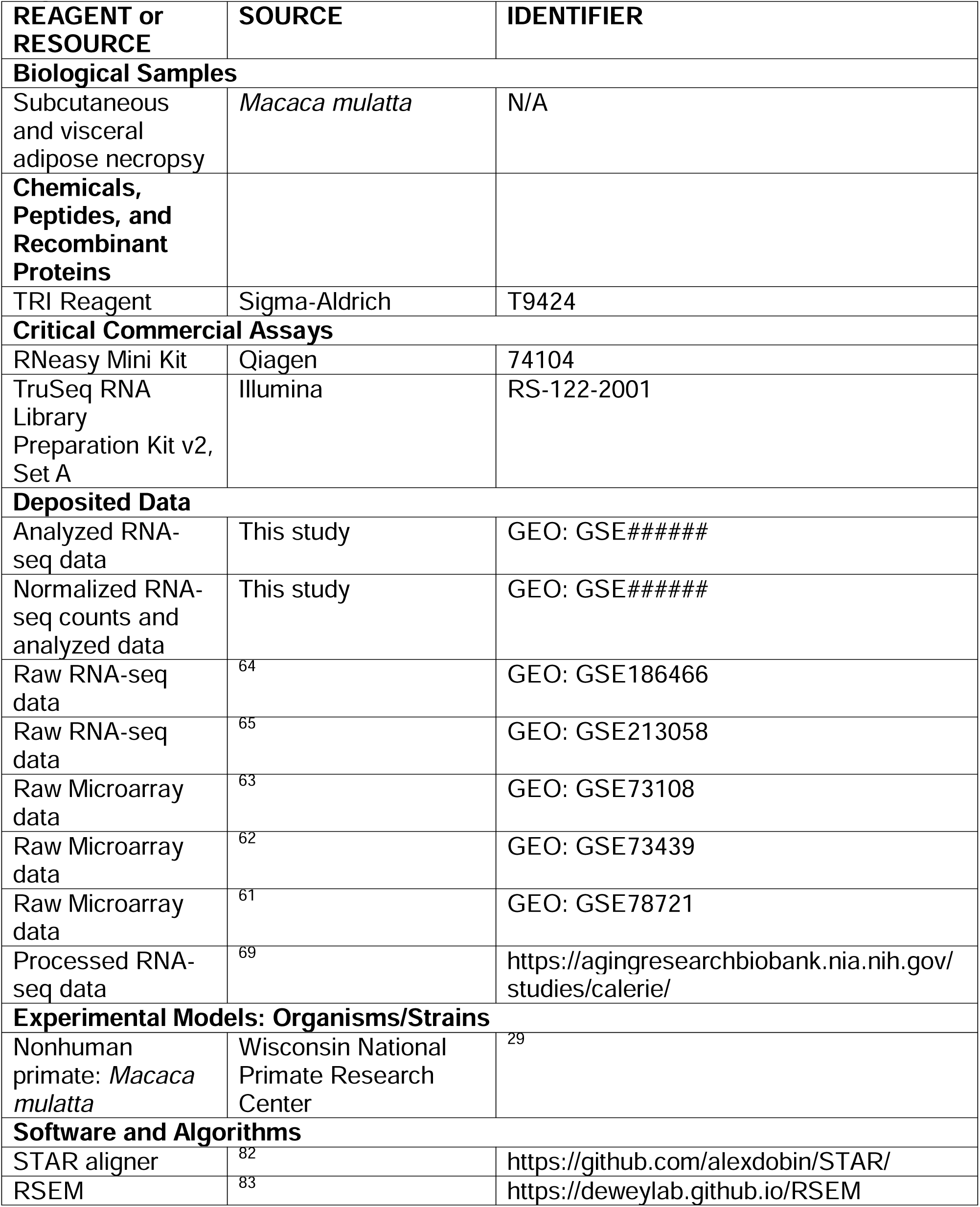
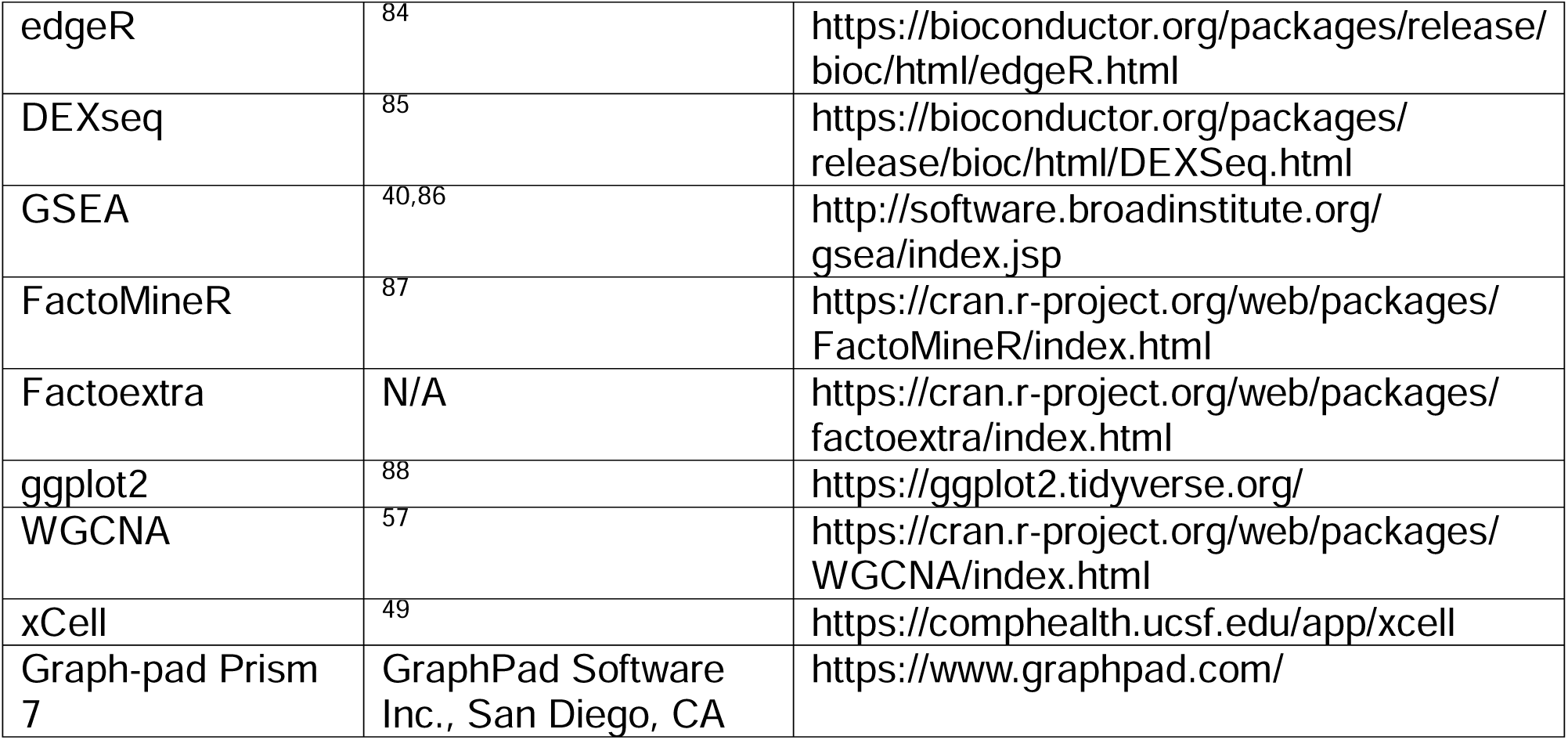

## Lead Contact and Materials Availability

Further information and request for resources and reagents should be directed to the Lead Contact, Rozalyn Anderson.

This study did not generate new unique reagents.

## Experimental Model and Subject Details

Animals were maintained and all procedures were performed at the Wisconsin National Primate Research Center (WNPRC) according to the guidelines for the ethical care and treatment of animals under approved protocols from the Institutional Animal Care and Use Committee of the Graduate School of the University of Wisconsin-Madison. Adult rhesus monkeys (*Macaca mulatta*) were enrolled in the study between 8-14 years of age and were fed a semi-purified, nutritionally-fortified, low-fat diet containing 15% protein and 10% fat. Upon initialization of the study, animals were randomized to Control (*ad libitum* fed) or CR diets. The CR diet was gradually implemented to 30% restriction based on individual baseline food intake. For the purpose of this study, animal assessments include body weight, body composition (via dual energy X-ray absorptiometry) and blood draws were collected for each animal within 6 months of necropsy.

## Method Details

### Biometrics

Body weight of each individual was assessed throughout the duration of the study. Total body fat mass, total body lean mass, total percent fat, appendicular lean mass, and total abdominal percent fat mass were assessed using whole body Dual Energy X-ray Absorptiometry (DEXA; model DPX-L, GE/Lunar Corp., Madison, WI) as previously described previously ^89^. Body weight, percentage fat and lean, and other biometric measures are reported as most recent measure before time of death. Insulin sensitivity was determined via the frequently sampled intravenous glucose tolerance test (FSIVGTT) ^90,91^.

### Blood chemistry

Complete blood counts were measured by standard laboratory procedures (Clinical Pathology Laboratory of the WNPRC). Other serum variables were measured as part of a serum chemistry panel (General Medical Laboratories, Madison, WI).

### Necropsy samples

Necropsy-obtained adipose tissue was collected from abdominal subcutaneous fat, and intra-abdominal visceral fat. Adipose tissue was stored at -80°C until samples were processed for transcriptomics.

### Transcriptomics

Greater than 300 mg of each adipose sample was homogenized in 1.5 mL of TRI Reagent (Sigma #T9424). RNA was then purified using the RNeasy Mini Kit (Qiagen #74104), and purified total RNA was then used to generate mRNA-sequencing libraries using the TruSeq RNA Library Preparation Kit v2, Set A (Illumina #RS-122-2001). Libraries were then run 2×100bp on an Illumina HiSeq2000.

### Analysis of sequencing data

The sequencing reads were trimmed using the trimming program Skewer (v0.1.123) ^92^. Trimmed reads were aligned to the *Macaca mulatta* reference genome ^93^ using STAR (v2.5.0a\n) ^82^. Read counts for each gene and isoform were quantified by RSEM ^83^. Differential expression analysis was done using edgeR (v 4.0.16)^84^, and the results used for pathway analysis by GSEA and ORA using KEGG (with the parameter organism = ‘hsa’).

### Software

Multifactor and principal component analysis performed with R package FactoMineR (http://factominer.free.fr/ CRAN) ^94^, with visualization done with Factoextra (CRAN). Cell type enrichment analysis from bulk RNA-seq expression data was performed using the xCell webtool ^49^(https://comphealth.ucsf.edu/app/xcell). Scaled cell type enrichment for each cell type-by-animal was calculated as the z-score of the raw xCell output of a given cell type across all cell types for that animal. Scaled cell type enrichment statistics were calculated using Student’s t-test. Additional data visualizations were performed using R package ggplot2 (CRAN).

### Weighted gene co-expression network analysis (WGCNA)

Using the WGCNA R package (v 1.73)^57^, clusters of expression profiles were identified. Using the transcript per million (TPM) value for every gene in all 16 samples (18,266 genes total), WGCNA identified and removed genes with too many missing values. Ultimately, clustering was run on 17,788 genes. For network topology analysis, we calculated the adjacencies using a soft thresholding power of 16 based on scale-free topology. We then transformed the adjacency into a topological overlap matrix (TOM), calculated the corresponding dissimilarity, and set a minimum module size to 30. Using the gene modules identified, KEGG pathway enrichment was calculated using over-representation analysis (ORA).

## Acknowledgments

We are grateful to Dr. Daniel Belsky for sharing transcriptional data from the CALERIE study ahead of publication. This work was supported by NIH AG040178, NIH AG074503, NIH training fellowships AG000213 (JPC). This work is supported by in part by NCRR/ORIP grants P51RR000167/P51OD011106 to the Wisconsin National Primate Research Center, University of Wisconsin-Madison, WI. This work was supported by the National Institutes of Health (P01AG011915, R01AG040178). This study was conducted using resources and facilities at the William S. Middleton Memorial Veterans Hospital, Madison, WI. The authors thank the University of Wisconsin Carbone Cancer Center Cancer Informatics Shared Resource, supported by P30 CA014520 for use of its services.

## Conflict of Interest

The authors declare no conflict of interest.

---

**Supplementary Figure S1:**
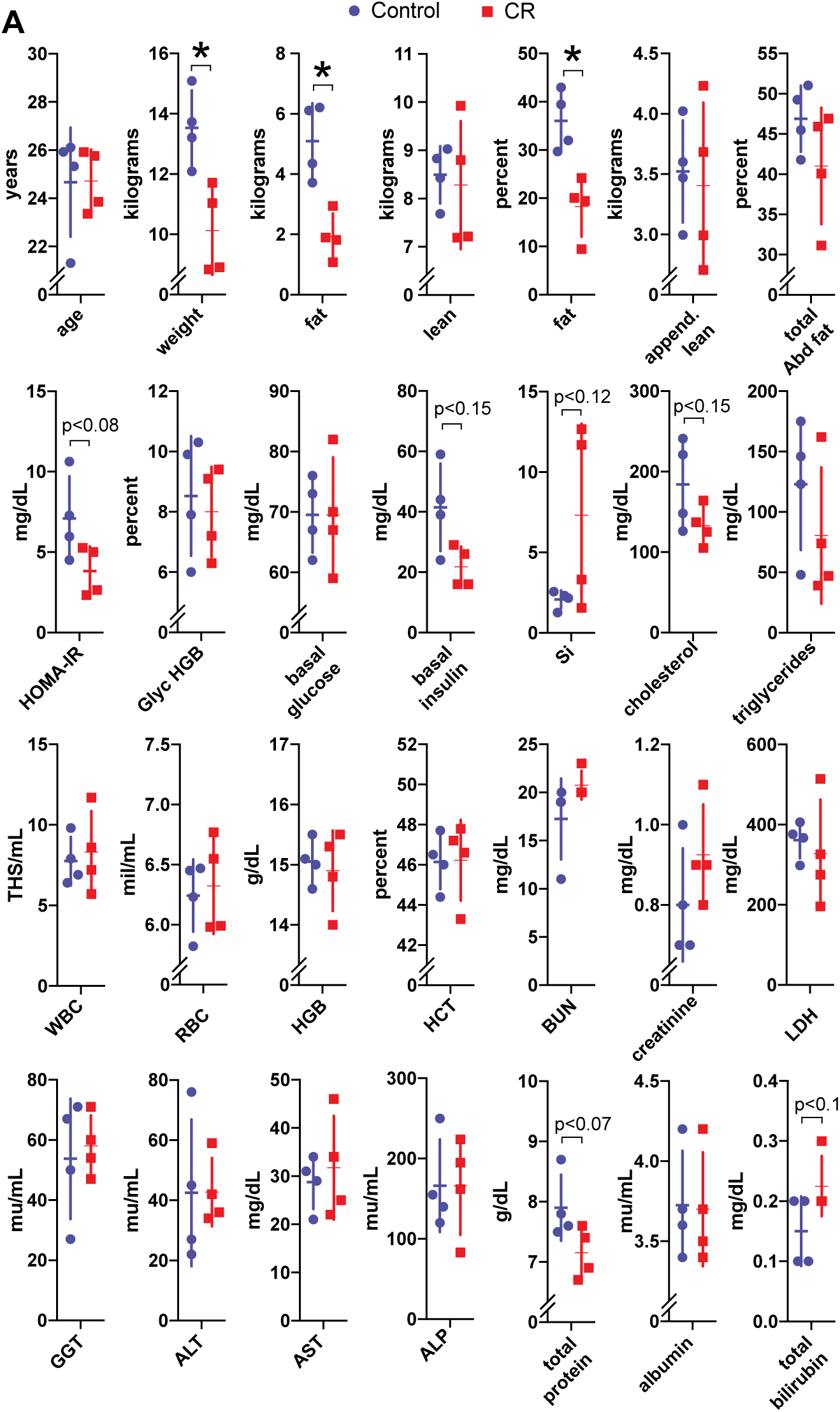
Blood analyses and biometric data for individuals on study. (**A**) Graphs showing biometric and blood chemistry measures for individuals on the Control or CR diet (n=4; bar, mean; line, SD). Significance indicated by * (p<0.05) or p-value listed from student t-test. Measures include: age, weight, fat, lean, appendicular lean mass, total and total abdominal fat percent, HOMA-IR, glycosylated-hemoglobin (Glyc HGB), basal glucose and insulin, insulin sensitivity (Si), cholesterol, triglycerides, white and red blood cell counts (WBC and RBC), hemoglobin (Hgb), hematocrit (Hct), blood urea nitrogen (BUN), creatinine, lactate dehydrogenase (LDH), gamma-glutamyl transferase (GGT), alanine aminotransferase (ALT), aspartate aminotransferase (AST), alkaline phosphatase (ALP), total protein, albumin, total bilirubin.

**Supplementary Figure S2:**
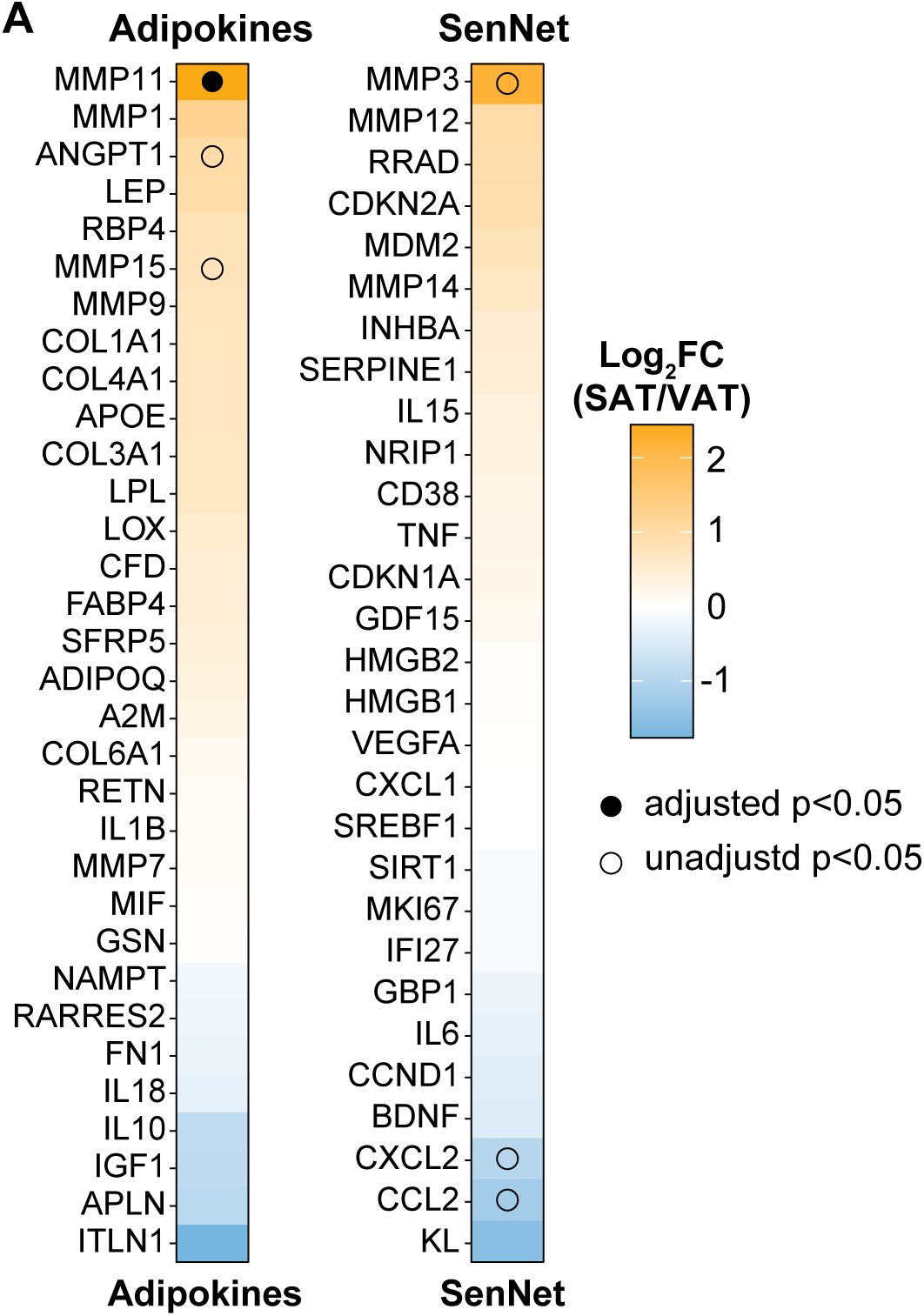
Depot-specific expression of Adipokines and SenNet factors. Heatmaps of SAT/VAT Log2FC for Adipokines (left) and SenNet factors (right). Filled dot denotes passing adjusted p<0.05, empty dot denotes unadjusted p<0.05. Related to Figure 1.

**Supplementary Figure S3:**
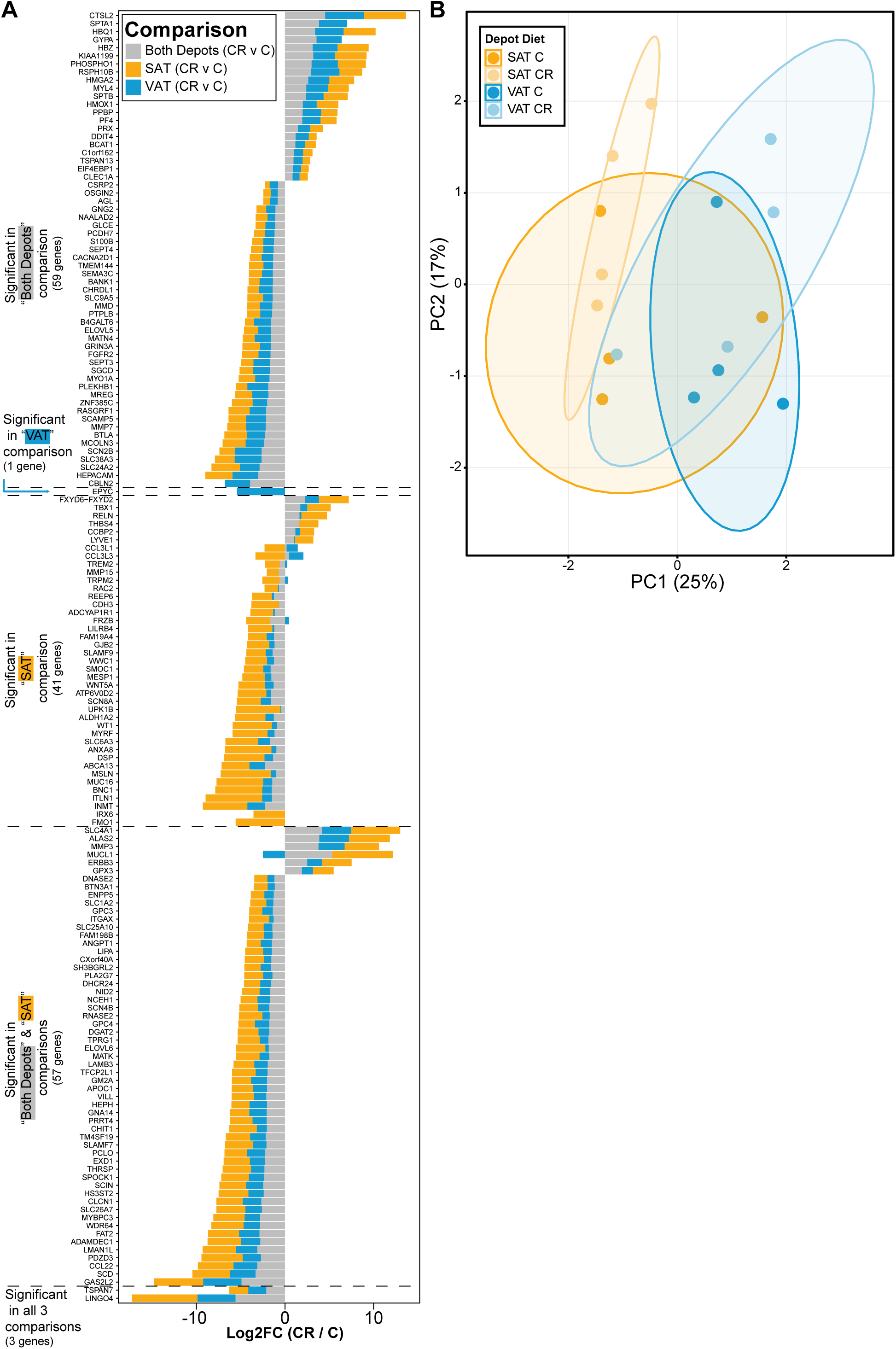
Depot-specific transcriptional response to CR. (**A**) Bar chart depicting the Log2FC (CR/C) of DEGs that are shared or unique to the three comparisons: both depots (gray), SAT (yellow) or VAT (blue). (**B**) Principal component analysis plot of the transcriptome for each group (n=4). Ellipse denotes 80% confidence level. Related to Figure 2.

**Supplementary Figure S4:**
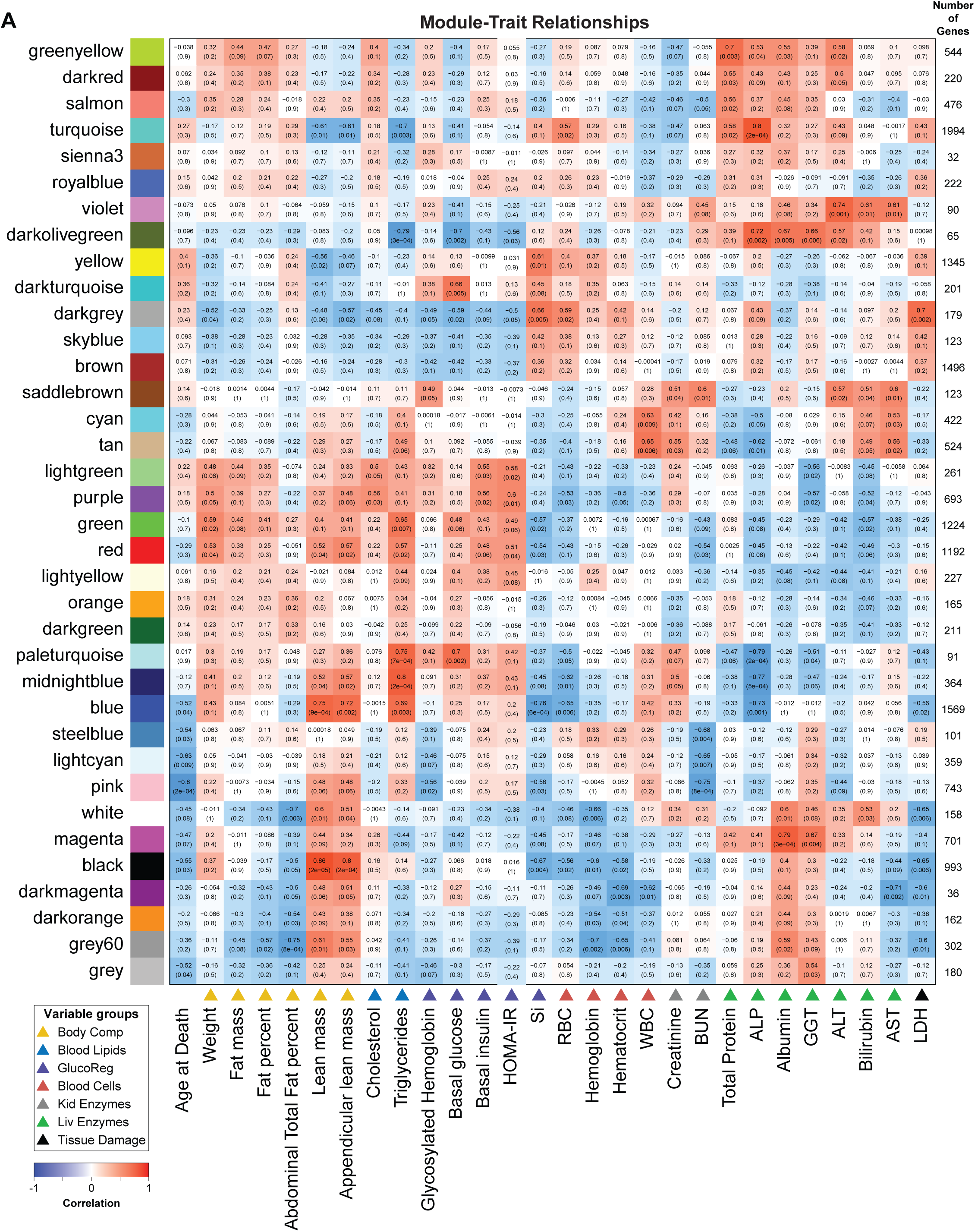
WGCNA module-trait associations. Correlation of module eigengenes with individual biometric measures (traits). Boxes contain module-trait correlations (top) and pvalue (bottom). Numbers of genes per module are listed on the right. Related to Figure 4.

**Supplementary Figure S5:**
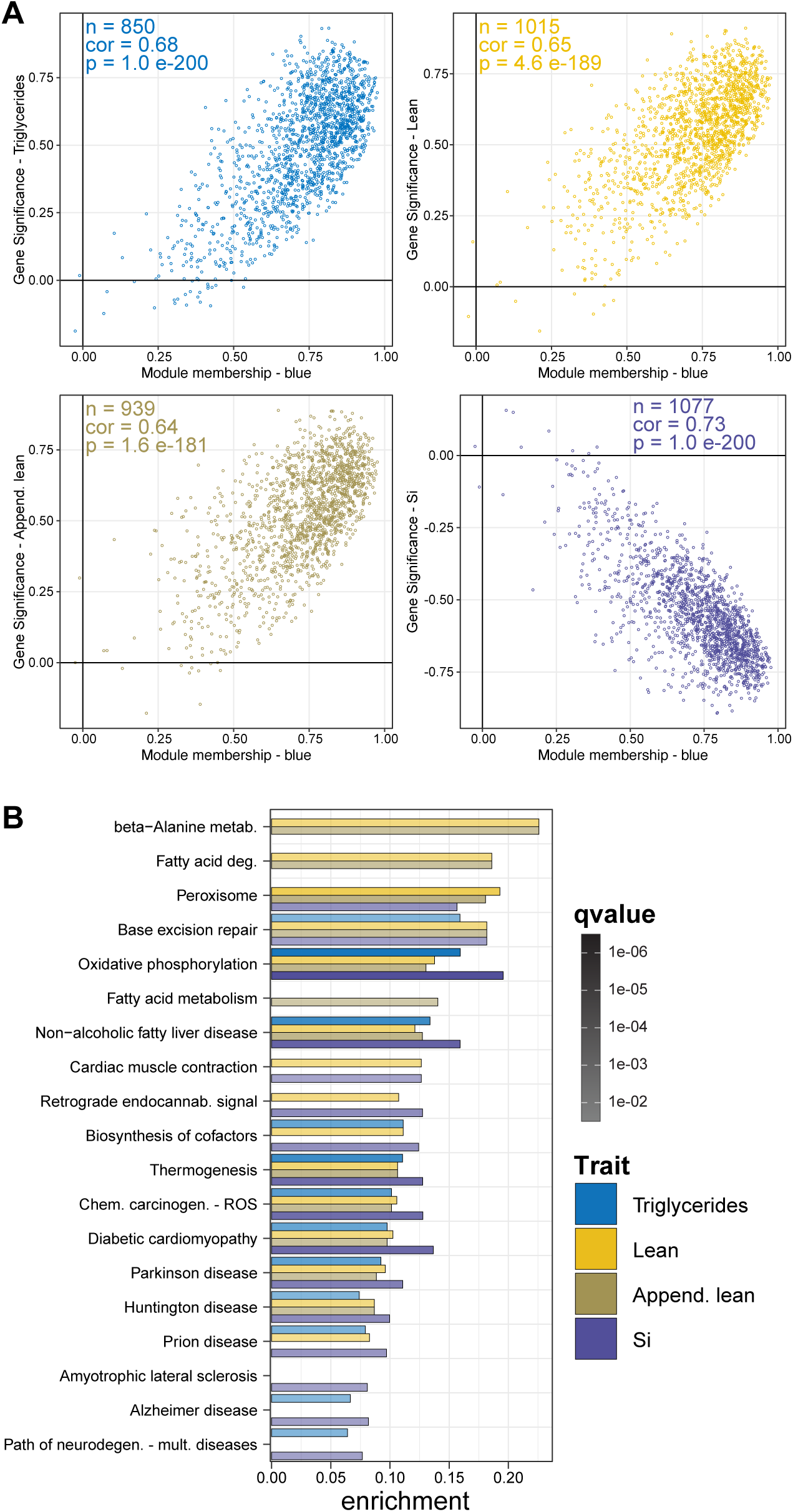
Blue module trait associations enrich for oxidative phosphorylation gene containing pathways. (**A**) Scatter plots of gene significance versus blue module-membership for genes associated with triglycerides, lean and appendicular mass, and insulin sensitivity (Si). n (number of genes), cor (correlation), p (pvalue); (**B**) Bar plot of KEGG pathways enriched via ORA (qvalue<0.05) for significant blue module genes (gene significance p<0.05) for each trait association. Related to Figure 5.

## Notes

### Competing Interest Statement

The authors have declared no competing interest.

